# Understanding the neurocognitive impact of outdoor PM10 and PM2.5 exposure: an in silico dosimetric modeling study using MPPD

**DOI:** 10.64898/2026.03.23.713644

**Authors:** Diego Ruiz-Sobremazas, Blanca Cativiela-Campos, María Cadalso, Angel Barrasa, Pilar Catalán-Edo, Cristian Perez-Fernandez, Beatriz Ferrer, Fernando Sánchez-Santed, Teresa Colomina, Caridad López-Granero

**Affiliations:** University of Zaragoza, Department of Psychology and Sociology, Teruel, Spain; Current: Department of Human Anatomy and Psychobiology, University of Murcia, Murcia, Spain; Department of Psychology and CIBIS, University of Almeria, Almeria 04120, Spain; Escuela Universitaria de Enfermería de Teruel (UNIZAR), Teruel, Spain; Department of Health Sciences, University of Burgos, Burgos 09001, Spain; Albert Einstein College of Medicine, Molecular Pharmacology Department, Bronx, NY, 10461, USA; Universitat Rovira I Virgili, Research Group in Neurobehavior and Health (NEUROLAB), Tarragona, Spain; Universitat Rovira I Virgili, Department of Psychology and Research Center for Behavior Assessment (CRAMC), Tarragona, Spain; Universitar Rovira I Virgili, Center of Environmental, Food and Toxicological Technology (TECNATOX), Reus, Spain

**Keywords:** Environmental Toxicology, Particulate Matter, MPPD model, Attentional functioning, Mental health, Oxidative Stress

## Abstract

Air pollution has been increasingly linked to adverse neurodevelopmental and neurodegenerative outcomes. While experimental and preclinical studies suggest that exposure to particulate matter (PM), particularly during gestation, may disrupt cognitive development, the impact of short-term PM exposure on cognitive and behavioral functioning in healthy young populations remains insufficiently explored in Spain. Moreover, few studies have incorporated individualized dosimetry models to estimate exposure more accurately. This study included 186 healthy young adults (mean age = 20.4 years) recruited from three Spanish cities (Teruel, Almería, and Talavera) characterized by different pollution levels. Ambient fine and coarse PM concentrations were recorded 8, 15, and 30 days prior to psychological assessment. Instead of relying solely on raw in situ environmental measurements, individualized PM deposition was estimated using the Multiple-Path Particle Dosimetry Model (MPPD), allowing a more biologically meaningful exposure approximation. Psychological outcomes were assessed using validated questionnaires: DASS-21 (depression, anxiety, stress), BIS-11 (impulsivity), UCLA Loneliness Scale, and SWLS (life satisfaction). Behavioral performance was evaluated using computerized versions of the Attentional Network Task (ANT) and the Stroop Task. Blood NRF2 concentrations were analyzed as a biomarker potentially related to oxidative stress mechanisms. In situ data indicated that Talavera presented the highest pollution levels, followed by Almería and Teruel. Linear regression analyses showed that coarse PM exposure across 8-, 15-, and 30-day windows significantly predicted poorer Executive Control Index performance in the ANT. Additionally, 15-day coarse PM and 30-day fine PM exposure were associated with greater cognitive interference. Oxidative stress markers were significantly associated with PM exposure levels. These findings support emerging evidence that short-term PM exposure may negatively affect executive and attentional processes even in healthy young adults. Further longitudinal research incorporating individualized exposure modeling is warranted to clarify causal pathways and underlying biological mechanisms.

**Graphical Abstract:** 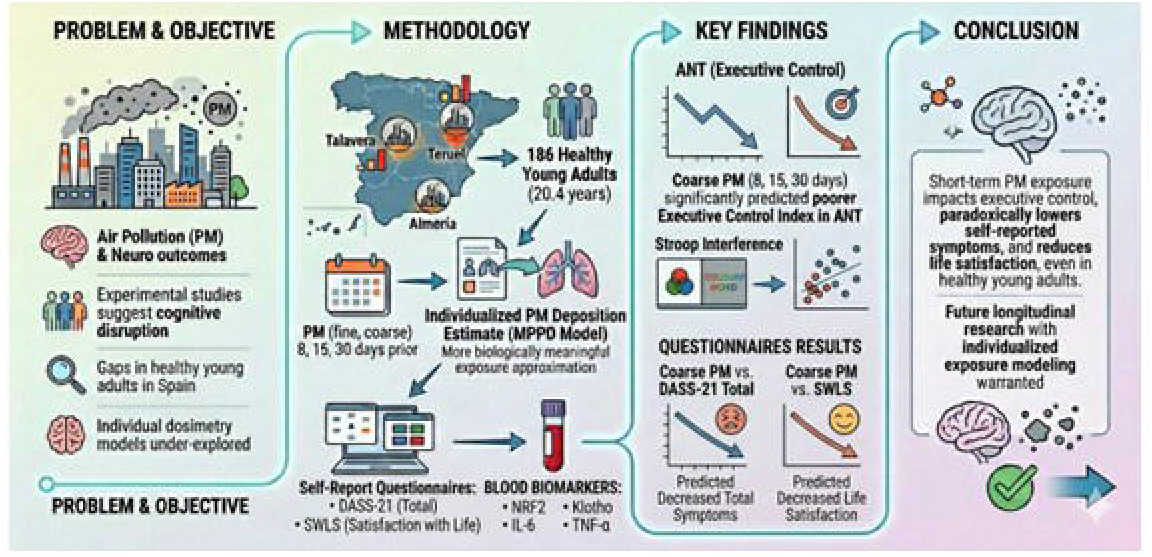

## 1. Introduction

Air pollution is widely recognized as one of the leading environmental risk factors contributing to global morbidity and mortality (Pozzer et al., 2023; Forastiere et al., 2024). Despite its broad recognition, there is no single, universally accepted definition of air pollution due to its intrinsic complexity. The World Health Organization (WHO, 2025) defines air pollution as the contamination of indoor or outdoor environments by chemical, physical, or biological agents that alter the natural characteristics of the atmosphere. Among the diverse components of air pollution, particulate matter (PM) has received particular attention because of its physicochemical properties and its capacity to penetrate biological systems (Onivefu & Imarhiagbe, 2024; National Institute of Environmental Health Sciences, 2025).

PM is typically classified according to aerodynamic diameter into coarse particles (PM_10_; ≤10 µm), fine particles (PM_2_._5_; ≤2.5 µm), and ultrafine particles (≤0.1 µm). Increasing evidence suggests that smaller particles may exert greater biological damage, as their reduced size facilitates deeper penetration into the respiratory tract and potential translocation into systemic circulation (Schraufnagel, 2020; Wyatt et al., 2020; Garcia et al., 2023). A substantial body of epidemiological research has examined the impact of PM exposure on respiratory and cardiovascular outcomes. Long-term cohort studies have consistently linked PM_2_._5_ exposure to an increased risk of chronic obstructive pulmonary disease, lower respiratory infections, and lung cancer (Kyung & Jeong, 2020). More recently, associations have been established between PM exposure and cardiovascular morbidity (Chanda et al., 2024), as well as neurodegenerative disorders (Cristaldi et al., 2022; Cativiela-Campos et al., 2025). These findings have progressively shifted attention toward the central nervous system (CNS) as a critical target of PM-related toxicity.

Early mechanistic hypotheses proposed that inhaled particles trigger systemic inflammatory responses through the release of pro-inflammatory mediators (Seaton et al., 1995). Subsequent research has identified multiple pathways through which PM may affect the CNS, including neuroinflammation, oxidative stress, disruption of the neurovascular unit, blood– brain barrier (BBB) dysfunction, and neuronal apoptosis (Calderón-Garciduenas et al., 2002, 2021; Cory-Slechta et al., 2023). Nevertheless, the extent to which these mechanisms translate into direct functional impairments in humans remains under active debate.

Two primary, non-mutually exclusive pathways have been proposed to explain PM-induced CNS alterations. The first involves a direct route, whereby ultrafine particles may translocate from the nasal epithelium to the brain via the olfactory bulb (Elder et al., 2006; Oberdöster et al., 2004; Graham et al., 2025). Additionally, fine particles deposited in the alveolar region may enter systemic circulation, facilitating the transport of soluble particle components that may compromise BBB integrity (Gunasingam et al., 2024; Calderón-Garciduenas et al., 2008). The second, indirect pathway suggests that PM exposure promotes systemic inflammation and oxidative stress, which in turn disrupt CNS homeostasis through immune activation and peripheral–central signaling cascades (Serafini et al., 2022; Rentschler & Kodavanti, 2024).

Preclinical evidence largely supports these mechanistic models. Experimental studies have demonstrated that PM exposure increases oxidative stress markers and neuroinflammatory responses (Zhang et al., 2023), induces apoptosis in CNS cells (Chang et al., 2019), reduces neurogenesis in animal models (Woodward et al., 2018), and is associated with structural brain alterations (Nephew et al., 2021; Ehsanifar et al., 2019). However, as highlighted by recent systematic reviews (Ruiz-Sobremazas et al., 2023; Rodulfo-Cardenas et al., 2023), there remains no clear consensus regarding the magnitude and specificity of PM effects on neurodevelopment, behavioral alterations, and global cognitive functioning.

Cognitive functioning encompasses a broad set of mental processes essential for adaptive daily behavior. According to the American Psychiatric Association (APA, 2018), cognitive functioning refers to the performance of mental processes such as perception, learning, memory, reasoning, judgment, and language. A growing number of preclinical studies have investigated the effects of PM exposure on specific cognitive domains, including memory (Ruiz-Sobremazas et al., 2025a; Hou et al., 2023; Fonken et al., 2012), learning (Ruiz-Sobremazas et al., 2025b; Patten et al., 2020), and social behavior (Weitekamp & Hofmann, 2021; Berg et al., 2020).

A similar trend is emerging in human research. Longitudinal cohort studies have explored associations between chronic PM exposure and cognitive aging (Qi et al., 2024; Wang et al., 2024; López-Granero et al., 2023), as well as cognitive development in school-aged children (Sunyer et al., 2015; Lett et al., 2017). Nevertheless, most human evidence remains observational in nature. To date, experimental research directly manipulating PM concentrations to evaluate acute behavioral or cognitive effects in humans is extremely limited. Notably, Faherty et al. (2025) conducted one of the first controlled studies modifying PM exposure levels and reported that higher PM concentrations impaired selective attention performance and reduced accuracy in emotion recognition tasks. Despite these advances, significant gaps remain regarding the causal relationship between PM exposure and specific cognitive domains in humans, as well as the mechanisms underlying these potential alterations. Further experimental research is therefore required to clarify the extent to which acute or short-term PM exposure directly influences cognitive functioning and behavioral outcomes.

One of the major challenges in environmental toxicology is the accurate assessment of the exposome. This requires the integration of in situ exposure measurements, which capture short-term and individual variability (Steinle et al., 2015), with in silico modeling approaches capable of simulating key environmental and exposure-related variables (Tsiros et al., 2022). The combination of these approaches improves exposure estimation and facilitates the identification of additional factors that may be influenced by varying PM concentrations (Houweling et al., 2024).

Based on previous experimental and epidemiological evidence, we hypothesized that higher PM exposure in three different exposure windows (8-, 15-, and 30-day exposure time) would be associated with poorer performance in attention and cognitive interference control tasks. Additionally, we incorporated systemic biological markers to explore whether the observed associations are more consistent with direct translocation mechanisms or indirect inflammatory pathways.

## 2. Methodology

### 2.1. Participants

One-hundredth and eighty-six Spanish-speaking participants from three cities (n = 120 from Teruel; n = 43 from Talavera; n = 23 from Almería), aged 18-39 (mean of 20.5 years old). Table 1 provides a summary of the demographic characteristics of the sample used. None of the cities are located close to each other, and are located at different regions of Spain: Teruel (in Aragón; 40°20⍰37⍰N 1°06⍰26⍰O), Almería (in Andalucía; ∼36°50⍰30⍰N 2°27⍰50⍰O) and Talavera de la Reina (Toledo; 39°58⍰00⍰N, 4°50⍰00⍰O). Exclusion criteria were: (i) not speaking Spanish, and (ii) diagnosis of intellectual disability or other psychiatry issues.

**Table 1.** Sample characteristics.

| Characteristics | Teruel ($n = 120$) | Almería ($n = 23$) | Talavera ($n = 43$) | Total | Comparisons<sup>1</sup> |
| --- | --- | --- | --- | --- | --- |
| Woman, $n$ (%) | 82 (68.3) | 18 (78.3) | 27 (62.8) | 127 (68.3) | $p = .441$ |
| Age, mean (SD) | 19.9 (2.4) | 22.6 (4.9) | 21.0 (2.4) | 20.4 (2.9) | $p = .005$ |
| Weight, mean (SD) | 65.3 (14.9) | 67.0 (19.3) | 64.7 (12.9) | 65.3 (14.4) | $p = .832$ |
| Height, mean (SD) | 165.1 (23.4) | 163.2 (7.7) | 169.3 (10.0) | 167.6 (9.6) | $p = .021$ |
| BMI, mean (SD) | 23.5 (6.5) | 24.8 (6.1) | 22.5 (3.6) | 23.7 (6.4) | $p = .315$ |
| Tidial Volume (SD) | 497.9 (71.3) | 466.8 (54.6) | 494.8 (59.1) | 493.3 (67.2) | $p = .125$ |
| Functional Residual Capacity<sup>2</sup> (SD) | 2.7 (0.5) | 2.5 (0.4) | 2.8 (0.5) | 2.7 (0.5) | $p = .073$ |
| Smoking, $n$ (%) | 14 (11.7) | 4 (17.4) | 7 (16.3) | 25 (13.4) | $p = .632$ |

<sup>1</sup> Due to the violation of Levene's test, all comparisons were performed with Welch correction.
<sup>2</sup> The functional residual capacity was calculated with the online Global Lung Function Initiative from the European Respiratory Society

Before assessment, participants gave written informed consent; participants did not receive any monetary compensation. The project was performed under the Declaration of Helsinki and approved by the local institutional ethics committees (reference number PI21-213) and in accordance with Regulation (EU) 2016/679, the General Data Protection Regulation, and with Organic Law 3/2018 on the Protection of Personal Data and Guarantee of Digital Rights (reference RAT 2023-284).

### 2.2. Real and physical data acquisition

The present cross-sectional study was conducted between March 2024 and May 2025. All data from participants (physiological, questionnaire, and neurocognitive outcomes) were obtained within only one session. First, participants were weighed and measured, after that, each participant completed all questionnaires, and after finishing those, they completed two computational tests (Attentional Network Task and Stroop). After finishing the computational tasks, participants were asked if they wanted to give blood for biochemical analysis.

### 2.3. Materials

#### 2.3.1. Assessment of mental health variables

Several questionnaires were used to screen different cognitive and mental health variables. First, we used a subjective measure of perceived health that consists of a Likert scale where participants must respond whether they considered that their own health is either bad or excellent. Also, we used the Spanish version of Satisfaction with Live Scale (SWLS; Pons et al., 2000).

The first variable analyzed is depression, anxiety and stress. We used the Spanish version of the Depression Anxiety and Stress Scale-21 (DASS-21; Lovibond & Lovibond, 1995). This test is composed of 21 items that screened three sub-dimensions: stress, anxiety and depression. Fonseca-Pedrero et al. (2010) showed good psychometric properties regarding Cronbach alpha for total symptoms (α =.90), for depression (α =.80), for anxiety (α =.73), and stress (α =.81). In addition, the Spanish DASS-21 version also achieved good punctuations in other psychometric outcomes.

The second variable analyzed is impulsivity. This variable was screened with the Spanish version of the Barratt Impulsiveness Scale (BIS-11; Patton, Stanford & Barratt, 1995). This questionnaire is composed of 30 items that screened total impulsiveness and three types of impulsivity: attentional, motor and non-planned impulsivity. The Spanish version of the BIS-11 was tested by Oquendo et al. (2001), revealing a good Cronbach alpha (α =.81).

The last emotional variable analyzed is loneliness. This variable was screened with the brief Spanish version of the UCLA loneliness scale. This questionnaire is composed of 10 items that screened loneliness. The questionnaire was tested by Velarde-Mayol et al. (2015), and obtained a good Cronbach alpha (α =.95).

#### 2.3.2. Assessment of neurocognitive variables

All tasks were programmed and administered using Psychopy (version 2023.2.3), an open-source software package for the creation and presentation of behavioral experiments. Psychopy allows precise control over stimulus presentation and response recording, ensuring millisecond accuracy in reaction time measurements. The experiment was conducted on a standard desktop computer and participants’ responses were recorded via keyboard input. We used a computerized version of the Attentional Networks Task (ANT) developed originally by Fan et al. (2002). This task involves participants responding to the direction of both cued (spatially or not) and uncued center arrow targets that are flanked by congruent, incongruent or neutral stimuli (see figure 1). The cue types measure how alerting or orienting attention networks are employed and the flanker types measure the ability to resolve conflict from visual stimuli. Participants, therefore, have to respond using the keyboard arrows (either right or left) to the direction of the central arrow that might be congruent or incongruent with the surrounding arrows. Furthermore, a control condition, where black squares were used as surrounding stimuli, was included within the procedure. We followed the same procedure explained in Fan et al. (2002) to calculate the data and the ratios for the orientation, alert, and cognitive functioning attentional networks using the following formulae: Orientation Index [(TRinvalid - TRvalid)*1000], Alert Index [(TRabsent - TRpresent)*1000], and Executive Function [(TRincongr - TRcongr)*1000]. All ratios were presented as reaction time in milliseconds (ms) as dependent variables to further analysis.

**Figure 1.**
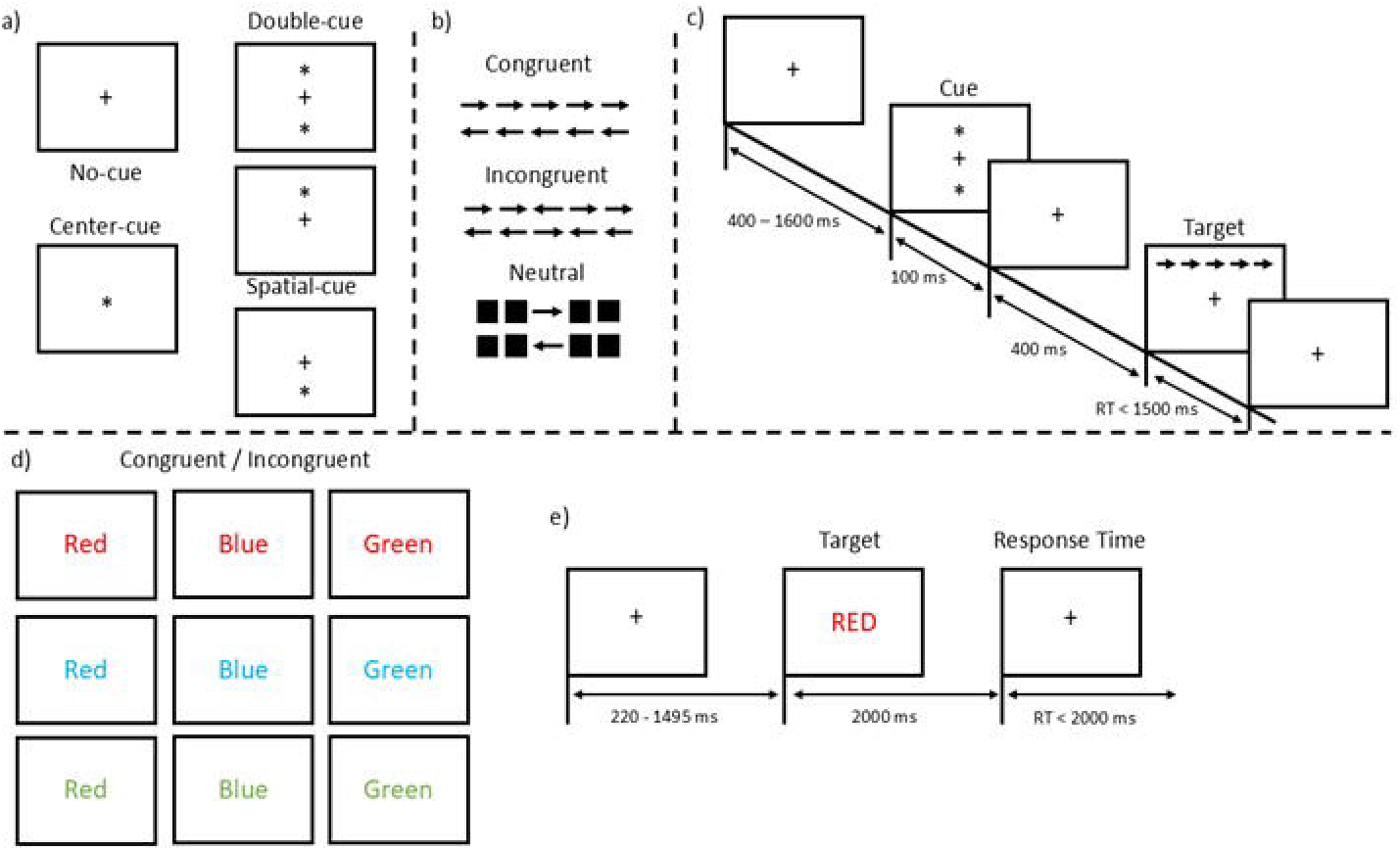
Graphical representation of ANT procedure (sections a, b and c). ln each trial, participants start by looking to a fixation point with a variable and random duration (400-1600 ms) until one of the cues depicted in image a) appears. After the cue, a fixation point reappears and, after 400ms, the target appears on the screen. Each participant needs to answer as soon as possible within 1500 ms or the trial is considered as an omission. The down part of the figure represents the STROOP procedure. Participants start the task with a fixation point with a random duration between 220-1495ms. After that, the target appears (one of the targets that appears in image d) and participants need to answer as soon as possible to the color of the word present (image e).

A computerized version of the Stroop Task, based on the original paradigm described by Stroop (1935), was administered. In this task, participants are required to respond to the font color of visually presented words while ignoring their semantic content, which can be congruent or incongruent with respect to the ink color. Congruent trials present color words whose meaning matches the displayed color (e.g., RED printed in red), whereas incongruent trials introduce a conflict between the word meaning and the ink color (e.g., RED printed in blue), thereby eliciting interference and engaging inhibitory control mechanisms. Participants respond via keyboard input, selecting the key corresponding to the ink color as quickly and accurately as possible. Stroop reaction time was used to calculate cognitive interference with the following formula: [(RTincongr/RTcongr)*100]

This procedure enables the quantification of cognitive interference and the efficiency of executive control processes involved in selective attention. All the tasks were performed using the same screens with the same characteristics (resolution of 1440 x 900, refreshment rate of 59.89 hz and 8-bits).

#### 2.3.3. In situ air pollution data

Air pollution information was taken from three different datasets. Those datasets are open-access from the Ministry for the Ecological Transition and the Demographic Challenge (for Almería), Regional Ministry of Sustainability, Environment and Blue Economy (for Talavera), and the Department of Environment and Tourism (for Teruel). Three different timelines were used to determine the number of molecules that were present in the air. Specifically, we selected 8-, 15-, and 30-days prior to collecting all the data. We selected these timelines because we wanted to differentiate between “acute” and “chronic” exposure to air pollutants.

Therefore, the air pollution data was taken from three different collectors: Almería (Code: ES1393A; 36.84133 altitude, -2.44672 longitude, 51 elevation above sea), Talavera (Code: ES2136A; 39.9638 altitude, -4.8257 longitude, 380 elevation above sea); Teruel (Code: ES1421A; 40.33639 altitude, -1.10667 longitude, 915 elevation above sea).

To obtain the final in situ PM concentration, each subject’s test day was considered. PM concentrations from the monitoring stations located in each city were calculated using the official data provided by governmental agencies. Finally, PM concentrations were averaged for each exposure period (8-, 15-, and 30-days).

#### 2.3.4. In silico air pathway simulation (MPPD)

To obtain a more accurate and individualized estimate of exposure to airborne contaminants, we employed the MPPD-v3.04 (Applied Research Associates, 2025), model to simulate the deposition of inhaled particles across the human respiratory tract (Anjilvel & Asgharian, 1995; Miller et al., 2016). Traditional exposure metrics based solely on ambient concentrations (e.g., µg/m^3^ measured at fixed-site monitoring stations) provide valuable information about environmental pollution levels but do not capture the substantial interindividual variability in inhalation, deposition, and clearance processes. As a result, relying exclusively on raw concentration data can lead to misclassification of personal exposure and an incomplete understanding of the biological dose that ultimately reaches target tissues.

The MPPD model offers a mechanistic framework that integrates particle characteristics (e.g., size distribution, density, hygroscopicity) with physiological parameters such as airway geometry, breathing patterns, and ventilation rates (Anjilvel & Asgharian, 1995; Miller et al., 2016; Mori, Ito & Sekine, 2024; Bui et al., 2020; Manojkumar et al., 2019). By simulating the transport and deposition of particles throughout the extrathoracic, tracheobronchial, and alveolar regions, the model provides a biologically meaningful estimate of the internal dose that is more closely aligned with toxicological relevance than ambient concentration alone. This is particularly important for contaminants whose health effects depend on the fraction of particles that penetrate deeply into the lungs or accumulate in specific anatomical regions.

Furthermore, the use of MPPD allows us to incorporate individualized or population-specific physiological inputs, thereby improving the ecological validity of exposure estimates. Factors such as age, sex, body size, and breathing mode (nasal vs. oral) can substantially influence deposition efficiency, and the model enables these sources of variability to be explicitly represented. This level of personalization is not achievable when using environmental concentration data alone, which assumes uniform exposure across individuals regardless of their physiological or behavioral characteristics.

By integrating environmental measurements with dosimetric modeling, our approach bridges the gap between external exposure and internal dose. This is essential for interpreting potential health effects, as the biological response to airborne contaminants is driven not by the concentration present in the environment but by the amount of material that actually deposits in the respiratory system. The MPPD model therefore provides a more refined and mechanistically grounded estimate of exposure, enhancing the accuracy of risk assessment and strengthening the link between environmental pollution and individual health outcomes. Table 2 summarized the specific variables to calculate individual data.

**Table 2.**
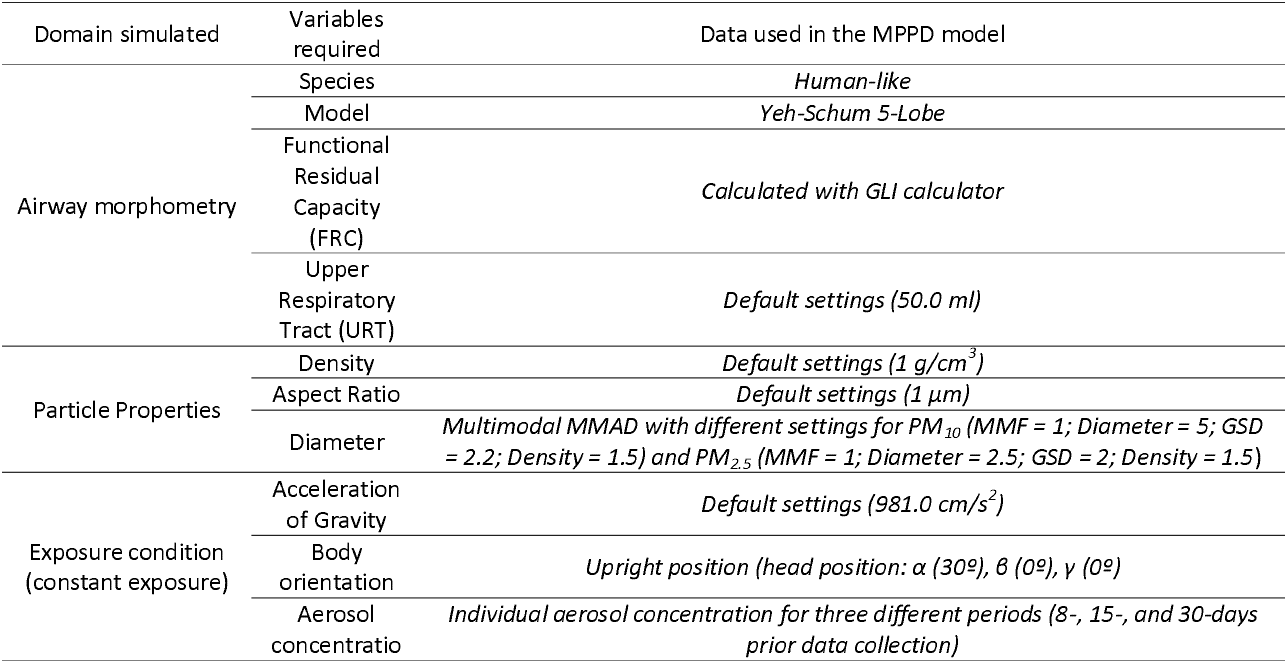
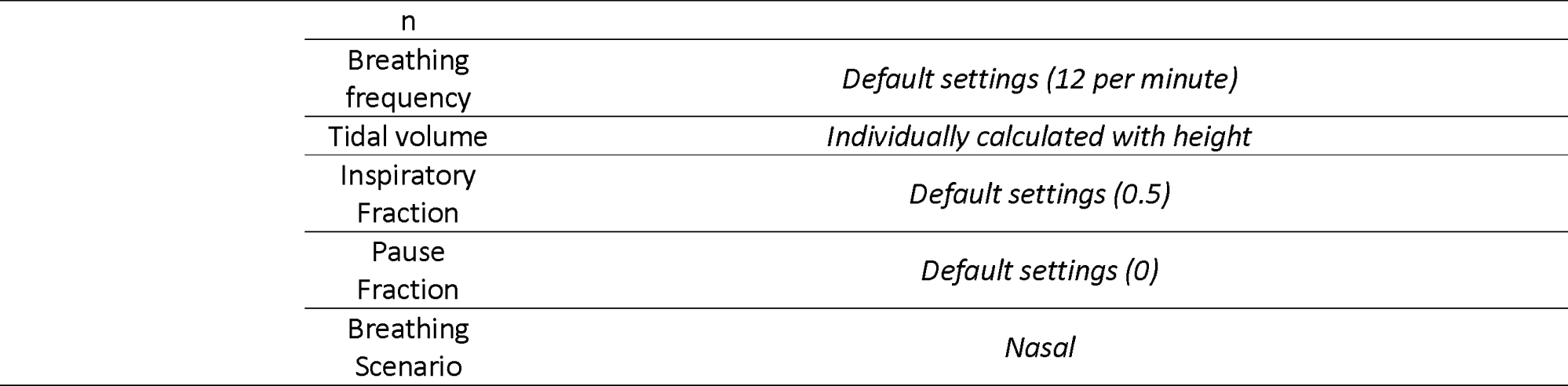
Variables required for the MPPD model and specific settings used for the calculations.

#### 2.3.5. Biochemical analysis

Blood samples were collected by specialized nursing staff at the same facilities where the research procedures were conducted. Due to funding constraints, biological samples could not be collected from participants in Talavera; therefore, only biological data from Teruel and Almería are included in this study. Whole blood was collected using Vacutainer® safety-lock blood collection systems (Vacutainer; Ref: 368652). Plasma samples were obtained in EDTA-coated tubes (Fisher Scientific; Ref: 12987666), while serum samples were collected in serum tubes (Fisher Scientific; BD Vacutainer SST II Advance; Ref: 10829270).

For plasma processing, EDTA-treated blood samples were stored at 4°C and processed within 1 hour of collection. Samples were centrifuged at 1700 × g for 10 minutes at 6°C in a refrigerated centrifuge. The plasma was aliquoted after centrifugation. For serum processing, samples were stored at 4°C under the same pre-processing conditions as plasma samples. Samples were centrifuged at 1.700 × g for 10 minutes at room temperature. No standardized clotting time at room temperature was applied, as samples were not centrifuged immediately after collection.

Following centrifugation, both plasma and serum samples were aliquoted and stored at −80°C until analysis. Samples were subjected to a maximum of two freeze–thaw cycles.

##### 2.3.5.1. ELISAs

Plasma and serum samples were analyzed using enzyme-linked immunosorbent assay (ELISA) kits for the quantification of the following biomarkers: interleukin-6 (IL-6; Invitrogen, Ref: EH2IL6), tumor necrosis factor alpha (TNF-α; Invitrogen, Ref: KAC1751), nuclear factor erythroid 2–related factor 2 (NRF2; Abcam, Ref: ab277397), and Klotho protein (Thermo Fisher Scientific, Ref: EEL200). All assays were performed according to the manufacturers’ instructions. All samples were analyzed in duplicate. Only measurements with a coefficient of variation below 15% between duplicates were included in the final analysis.

Standard curves were generated for each plate according to the manufacturer’s recommendations, and concentrations were calculated using a four-parameter logistic regression model. The lower limit of detection (LLOD) for each assay was as specified by the manufacturer.

A pilot analysis performed in EDTA-plasma samples yielded cytokine concentrations below the LLOD for all analytes. Consequently, all subsequent ELISA measurements were conducted in serum samples. Given that the study population did not present clinically overt inflammatory conditions, low circulating levels of pro-inflammatory cytokines were anticipated, in line with previous literature (Kleiner et al., 2013). All ELISAs were read using Infinite® F50 absorbance reader (TECAN, Ref: 30190077) following each ELISA’s instruction.

### 2.4. Data Analysis

Two different statistical analyses were performed with the obtained data. First, environmental and MPPD simulated data were compared using two-way ANOVAs where PM and MPPD simulated outcomes between cities are compared. Previously performing this analysis, equality of variances was tested with Levene’s test. If the test shows no equal variances, therefore a Welch ANOVA will be performed because it works perfectly with unequal variances (Welch, 1951; Zimmerman, 2004) and different sample sizes (Zimmerman, 2004). Spearman correlations are performed with the data to determine the relationship between variables.

To examine the association between environmental pollution and the behavioral outcomes of interest, linear regression models were fitted with simulated particle absorption (percentage absorbed simulated by the MPPD model multiplied by the real PM concentration) as the main predictor. PM absorption was derived from the MPPD model and was used as a continuous indicator of contextual environmental pollution load, reflecting differences in ambient exposure across areas with varying levels of air pollution rather than precise individual-level exposure. Because the primary variability in simulated absorption occurs between areas with different pollution loads, and not within each specific area, city was not included as a covariate neither as a factor in the main regression models, in order to avoid over-adjustment that would remove the contextual variability of interest. Accordingly, regression coefficients are interpreted as associations at the environmental-contextual level, and not as individual causal effects. Same analysis is performed with the 8-, 15-, and 30-exposure period time. Sex was included as covariable in all the regression analyses.

Given the moderate sample size and the subtle effect sizes typically observed in environmental neurotoxicology studies, regression models were intentionally kept parsimonious to reduce the risk of overfitting and unstable parameter estimates. Therefore, the primary analyses focused on the association between contextual PM absorption estimates and behavioral outcomes, rather than attempting to adjust for a large number of potential environmental covariates.

Separate regression models were estimated for each behavioral outcome. Assumptions of linearity, homoscedasticity, and normality of residuals were assessed through visual inspection of residual plots and summary statistics, with no major violations detected. Statistical significance was set at p <.05. All analyses were conducted using JASP 0.19.3.0 and GraphPad Prism 8.

## 3. Results

### 3.1. Sample characteristics and sociodemographic information

All participants were recruited from the University of Zaragoza at the campus of Teruel, the University of Almería, and people living in the city of Talavera. No differences were detected in the percentage of women that participated in the study, nevertheless, females correspond to 68.3% of the total sample; furthermore, other variables linked to the MPPD model and to other demographic aspects were not significant. We found significant differences in age (F_(2, 47.119)_ = 6.072; p =.005; η^2^_p_ =.098). Bonferroni post-hoc analysis revealed significant differences between Teruel and Almería (t = -4.246; p <.001), while the rest comparisons did not reach significant levels (p ≈.083). Also, a significant difference was detected in height (F_(2, 56.868)_ = 4.165; p =.021; η^2^_p_ =.034). Post-hoc comparisons revealed that Almería and Talavera had different heights (t = -2.489; p =.041), while the rest did not reach significant differences. No other comparisons revealed significant differences (Table 1).

### 3.2. In situ PM_10_ and PM_2.5_ concentration and MPPD simulation outcomes

PM10 and PM2.5 concentrations were analyzed in three different time periods previous data collection (8-, 15-, and 30-days). Results show that we, therefore, selected different participants which are exposed to different concentrations (p <.001) across all exposure time windows. Nevertheless, the air pollution dynamics was not stable across all comparisons, revealing different dynamics in PM10 and PM 2.5 concentrations across the cities (Table 2).

In terms of PM2.5 concentrations, Talavera achieved higher concentrations than Teruel, and Teruel higher from Almería [8-days, (Talavera, M = 21.885 µg/m^3^, SD = 7.005; Teruel, M = 9.927, SD = 6.930; Almería, M = 5.751, SD = 0.168); 15-days (Talavera, M = 15.531 µg/m^3^, SD = 3.209; Teruel, M = 9.899 µg/m^3^, SD = 5.485; Almería, M = 5.696 µg/m^3^, SD = 0.121); 30-days (Talavera, M = 16.054 µg/m^3^, SD = 0.465; Teruel, M = 9.036 µg/m^3^, SD = 2.039; Almería, M = 6.181 µg/m^3^, SD = 0.084)]. However, in terms of PM10 concentrations changed the city distribution, being the most polluted city Talavera followed by Almería and then Teruel [8-days (Talavera, M = 36.47 µg/m^3^, SD = 11.674; Almería, M = 21.46 µg/m^3^, SD = 0.858; Teruel, M = 14.20 µg/m^3^, SD = 10.901), 15-days (Talavera, M = 25.88 µg/m^3^, SD = 5.349; Almería, M = 21.15 µg/m^3^, SD = 0.844; Teruel, M = 14.41 µg/m^3^, SD = 8.617), 30-days (Talavera, M = 26.76 µg/m^3^, SD = 0.775; Almería, M = 24.54 µg/m^3^, SD = 0.496; Teruel, M = 13.61 µg/m^3^, SD = 3.197).

Results show that those PM concentrations differ between cities in all of the performed comparisons (p < 0.001). Nevertheless, the air pollution dynamics was not stable across all comparisons. Table 2 summarizes all the comparisons regarding time periods and cities in air pollution measures and Figure 2 shows graphically that dynamic.

**Figure 2.**
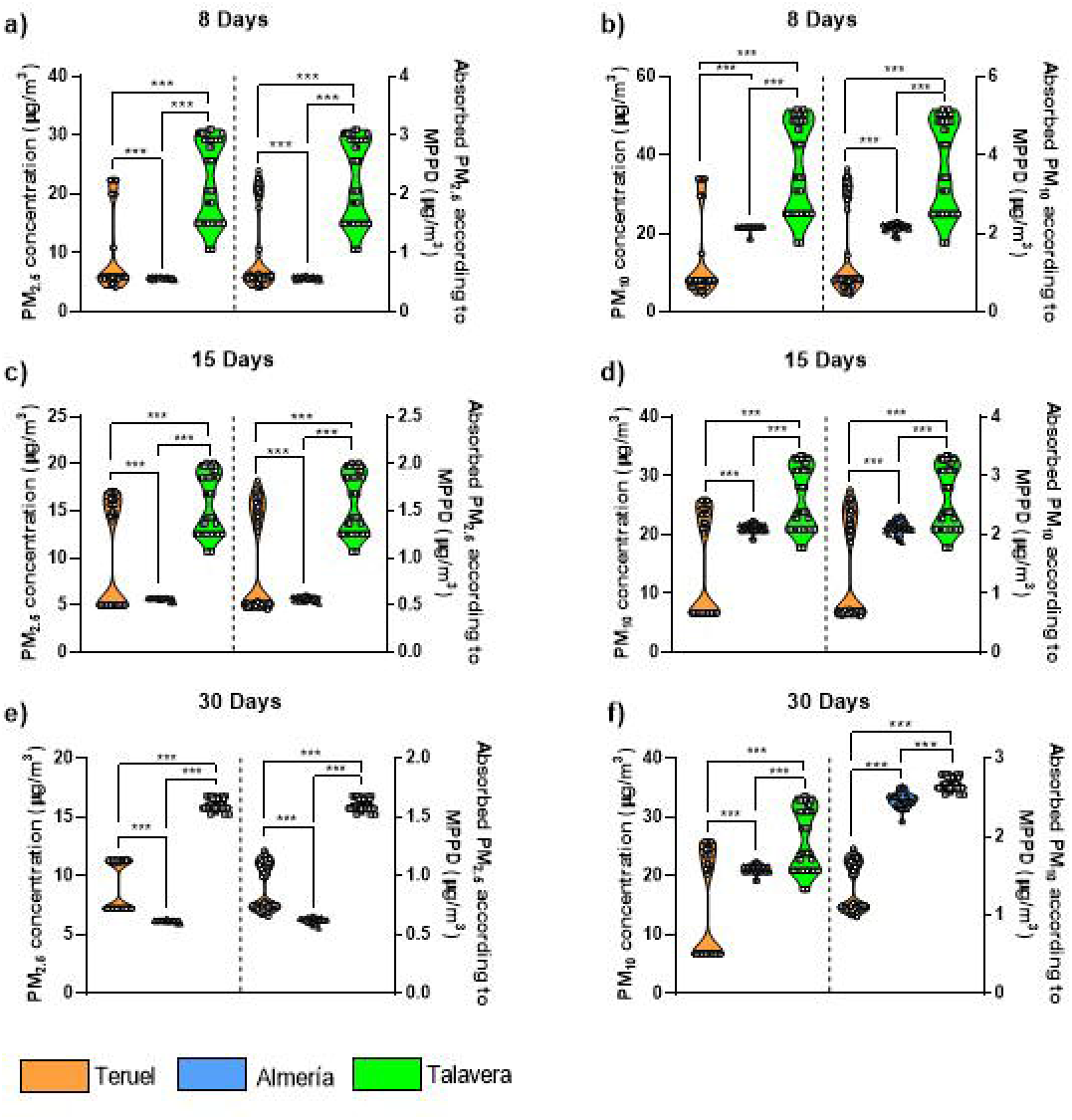
Graphical representation of the PM2.5 (graphs a, c, e) and PM10 (graphs b, d, f). Graphs a) and d) show PM concentration in 8 days; graphs c) and d) show PM concentration in 15 days; graphs e) and f) show PM concentration in 30 days. The left Y axis corresponds to *in* situ air pollution outcomes, while the right Y axis corresponds to absorbed PM according to the MPPD simulation *(in vitro)*. Each graph depicts individual data. ***corresponds to p < 0.001. Teruel (n = 120), Almería (n = 23), Talavera (n = 43).

As described earlier, the MPPD model was developed to improve the representation of air pollution exposure and to assess its effects across different medical endpoints. Nevertheless, this model personalizes the air pollution exposure. Therefore, the total alveolar deposit fraction was used to calculate *in vitro* PM exposures. Those exposures followed the same dynamics than *in situ* outcomes. Absorbed PM according to MPPD model is depicted in Figure 2. MPPD’s *in silico’s* simulation exposure also differ between cities as can be seen in Table 2. Talvera’s subjects absorbed more PM2.5 than Teruel and then Almería [8-days, (Talavera, M = 2.186 µg/m^3^, SD = 0.700; Teruel, M = 0.982 µg/m^3^, SD = 0.671; Almería, M = 0.579 µg/m^3^, SD = 0.026), Figure 2a; 15-days (Talavera, M = 1.551 µg/m^3^, SD = 0.321; Teruel, M = 0.975 µg/m^3^, SD = 0.526; Almería, M = 0.573 µg/m^3^, SD = 0.023), Figure 2c; 30-days (Talavera, M = 1.604 µg/m^3^, SD = 0.046; Teruel, M = 0.896 µg/m^3^, SD = 0.187; Almería, M = 0.622 µg/m^3^, SD = 0.023), Figure 2e]. Same as before, the pattern changed, being the subjects from Talavera those that absorbed more PM10 than Almería and then Teruel [8-days, (Talavera, M = 3.644 µg/m^3^, SD = 1.166; Teruel, M = 1.402 µg/m^3^, SD = 1.058; Almería, M = 2.159 µg/m^3^, SD = 0.112), Figure 2b; 15-days (Talavera, M = 2.586 µg/m^3^, SD = 0.534; Teruel, M = 1.417 µg/m^3^, SD = 0.828; Almería, M = 2.128 µg/m^3^, SD = 0.114), Figure 2d; 30-days (Talavera, M = 2.673 µg/m^3^, SD = 0.077; Teruel, M = 1.349 µg/m^3^, SD = 0.294; Almería, M = 2.469 µg/m^3^, SD = 0.101), Figure 2f].

**Table 2a.**
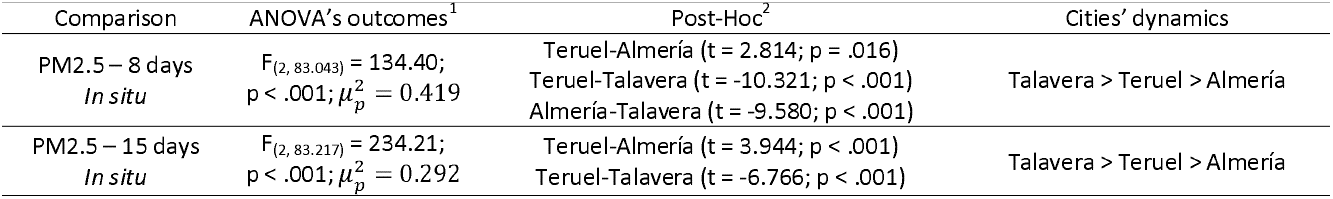
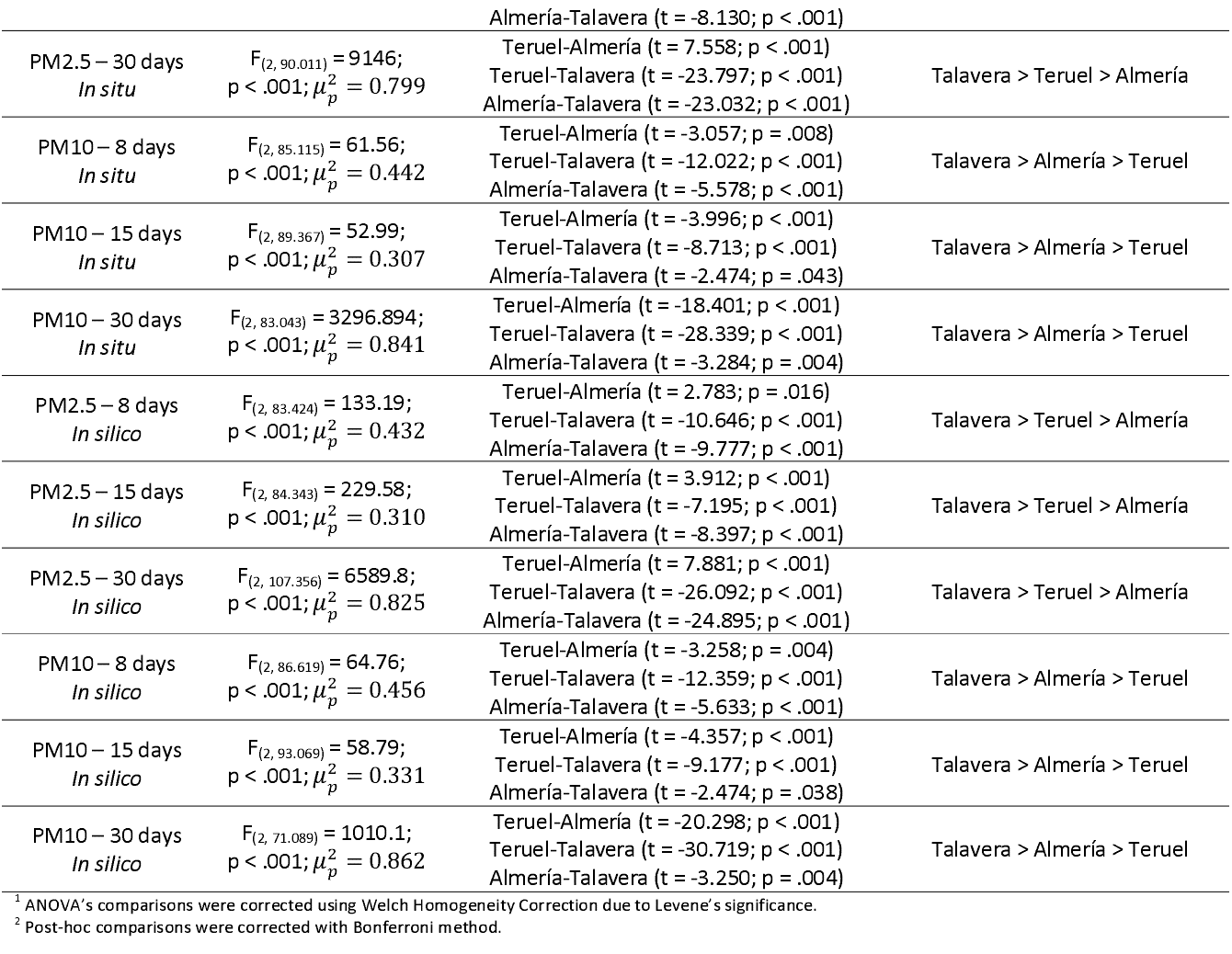
ANOVA’s comparisons from *in situ* and *in silico* PM_2.5_ and PM_10_ measures across different exposure time windows.

| Comparison | ANOVA's outcomes<sup>1</sup> | Post-Hoc<sup>2</sup> | Cities' dynamics |
| --- | --- | --- | --- |
| PM<sub>2.5</sub> – 8 days<br>*In situ* | $F_{(2, 83.043)} = 134.40$;<br>$p < .001$; $\mu_p^2 = 0.419$ | Teruel-Almería ($t = 2.814$; $p = .016$)<br>Teruel-Talavera ($t = -10.321$; $p < .001$)<br>Almería-Talavera ($t = -9.580$; $p < .001$) | Talavera > Teruel > Almería |
| PM<sub>2.5</sub> – 15 days<br>*In situ* | $F_{(2, 83.217)} = 234.21$;<br>$p < .001$; $\mu_p^2 = 0.292$ | Teruel-Almería ($t = 3.944$; $p < .001$)<br>Teruel-Talavera ($t = -6.766$; $p < .001$) | Talavera > Teruel > Almería |
|  |  | Almería-Talavera (t = -8.130; p < .001) |  |
| PM2.5 – 30 days<br>*In situ* | $F_{(2, 90.011)} = 9146;$<br>$p < .001; \mu_p^2 = 0.799$ | Teruel-Almería (t = 7.558; p < .001)<br>Teruel-Talavera (t = -23.797; p < .001)<br>Almería-Talavera (t = -23.032; p < .001) | Talavera > Teruel > Almería |
| PM10 – 8 days<br>*In situ* | $F_{(2, 85.115)} = 61.56;$<br>$p < .001; \mu_p^2 = 0.442$ | Teruel-Almería (t = -3.057; p = .008)<br>Teruel-Talavera (t = -12.022; p < .001)<br>Almería-Talavera (t = -5.578; p < .001) | Talavera > Almería > Teruel |
| PM10 – 15 days<br>*In situ* | $F_{(2, 89.367)} = 52.99;$<br>$p < .001; \mu_p^2 = 0.307$ | Teruel-Almería (t = -3.996; p < .001)<br>Teruel-Talavera (t = -8.713; p < .001)<br>Almería-Talavera (t = -2.474; p = .043) | Talavera > Almería > Teruel |
| PM10 – 30 days<br>*In situ* | $F_{(2, 83.043)} = 3296.894;$<br>$p < .001; \mu_p^2 = 0.841$ | Teruel-Almería (t = -18.401; p < .001)<br>Teruel-Talavera (t = -28.339; p < .001)<br>Almería-Talavera (t = -3.284; p = .004) | Talavera > Almería > Teruel |
| PM2.5 – 8 days<br>*In silico* | $F_{(2, 83.424)} = 133.19;$<br>$p < .001; \mu_p^2 = 0.432$ | Teruel-Almería (t = 2.783; p = .016)<br>Teruel-Talavera (t = -10.646; p < .001)<br>Almería-Talavera (t = -9.777; p < .001) | Talavera > Teruel > Almería |
| PM2.5 – 15 days<br>*In silico* | $F_{(2, 84.343)} = 229.58;$<br>$p < .001; \mu_p^2 = 0.310$ | Teruel-Almería (t = 3.912; p < .001)<br>Teruel-Talavera (t = -7.195; p < .001)<br>Almería-Talavera (t = -8.397; p < .001) | Talavera > Teruel > Almería |
| PM2.5 – 30 days<br>*In silico* | $F_{(2, 107.356)} = 6589.8;$<br>$p < .001; \mu_p^2 = 0.825$ | Teruel-Almería (t = 7.881; p < .001)<br>Teruel-Talavera (t = -26.092; p < .001)<br>Almería-Talavera (t = -24.895; p < .001) | Talavera > Teruel > Almería |
| PM10 – 8 days<br>*In silico* | $F_{(2, 86.619)} = 64.76;$<br>$p < .001; \mu_p^2 = 0.456$ | Teruel-Almería (t = -3.258; p = .004)<br>Teruel-Talavera (t = -12.359; p < .001)<br>Almería-Talavera (t = -5.633; p < .001) | Talavera > Almería > Teruel |
| PM10 – 15 days<br>*In silico* | $F_{(2, 93.069)} = 58.79;$<br>$p < .001; \mu_p^2 = 0.331$ | Teruel-Almería (t = -4.357; p < .001)<br>Teruel-Talavera (t = -9.177; p < .001)<br>Almería-Talavera (t = -2.474; p = .038) | Talavera > Almería > Teruel |
| PM10 – 30 days<br>*In silico* | $F_{(2, 71.089)} = 1010.1;$<br>$p < .001; \mu_p^2 = 0.862$ | Teruel-Almería (t = -20.298; p < .001)<br>Teruel-Talavera (t = -30.719; p < .001)<br>Almería-Talavera (t = -3.250; p = .004) | Talavera > Almería > Teruel |

<sup>1</sup> ANOVA's comparisons were corrected using Welch Homogeneity Correction due to Levene's significance.
<sup>2</sup> Post-hoc comparisons were corrected with Bonferroni method.

Furthermore, the MPPD model calculates several variables related to air pollution exposure. However, we only selected those variables that are directly related with the main objective of the present work. Between those variables, we selected deposit fractions across all the respiratory tract (alveolar, conducting airways, and head). Interestingly, the ANOVA revealed differences in total alveolar deposit fraction [F_(2, 57.142)_ = 7.698, p =.001; 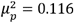, Figure 3a], where Talavera achieved lower total alveolar deposit fraction than Almería and Teruel [post-hoc comparisons: Teruel-Almería (t = -0.632, p_bonf_ = 1); Teruel-Talavera (t = 4.620, p_bonf_ <.001); Almería-Talavera (t = 3.735; p_bonf_ <.001). In addition, the total percentage deposit fraction in the conducting airways also revealed differences between cities [F_(2, 59.002)_ = 16.695, p <.001; 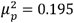, Figure 3b]. However, Talavera subjects had higher percentage deposit fraction in the conducting airways than Almería and Teruel [post-hoc: Teruel-Talavera (t = -1.706; p_bonf_ =.269), Teruel-Talavera (t = -6.655; p_bonf_ <.001), Almería-Talavera (t = -3.075; p_bonf_ =.007). The last variable analyzed is the total head deposition, revealing no differences (p =.128, Figure 3c).

**Figure 3.**
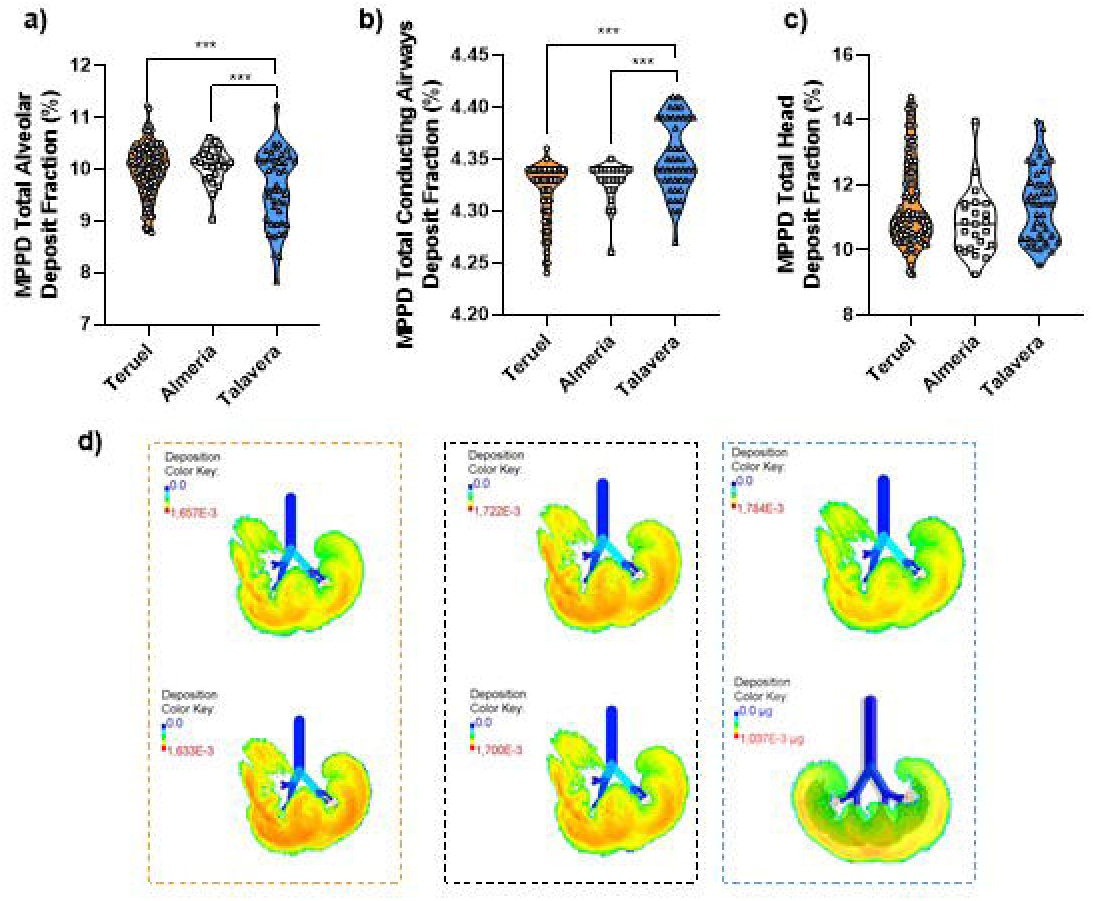
Graphical representation of main MPPD-outcomes related to the objective of the present work. The upper violin graphs depict total alveolar deposit fraction (%, image a), total conducting airways deposit fraction (%, image b), and total head deposit fraction (%, image c). Image d shows graphical simulations of a 5-lobe non-symmetric lung in Teruel (orange), Almería (black) and Talavera (blue), The lower lung graph in Talavera (blue) also depicts 10 lung ramifications. ***p <.001

### 3.3. Correlations between in silico PM10 and PM2.5 concentrations and cognitive/behavioral variables

Prior calculating linear regression, correlation analyses were performed between variables to determine the present relationships. Only the significant correlations are shown, therefore, those correlations that were not included did not achieve significant levels. Due to the no-normality of our data, non-parametric Spearman correlations were calculated.

As can be seen in Figure 4, the correlation matrix revealed different variable dynamics. First, several negative correlations were detected between PM2.5 exposure for 15-, and 30-days and emotional outcomes. Being exposed to PM2.5 for 15 days is negatively related with DASS-21 scores (depression, p =.020, ρ = -.171; anxiety, p =.033, ρ = -.157; stress, p =.005, ρ = -.208; and total punctuation, p =.006, ρ = -.203). The exposure window of 30-days also revealed the same negative correlations with DASS-21 (depression, p =.007, ρ = -.196; anxiety, p =.015, ρ = -.179; stress, p <.001, ρ = -.247; and total punctuation, p =.001, ρ = -.235). Higher PM concentrations are related with lower punctuations in the psychological test performed.

**Figure 4.**
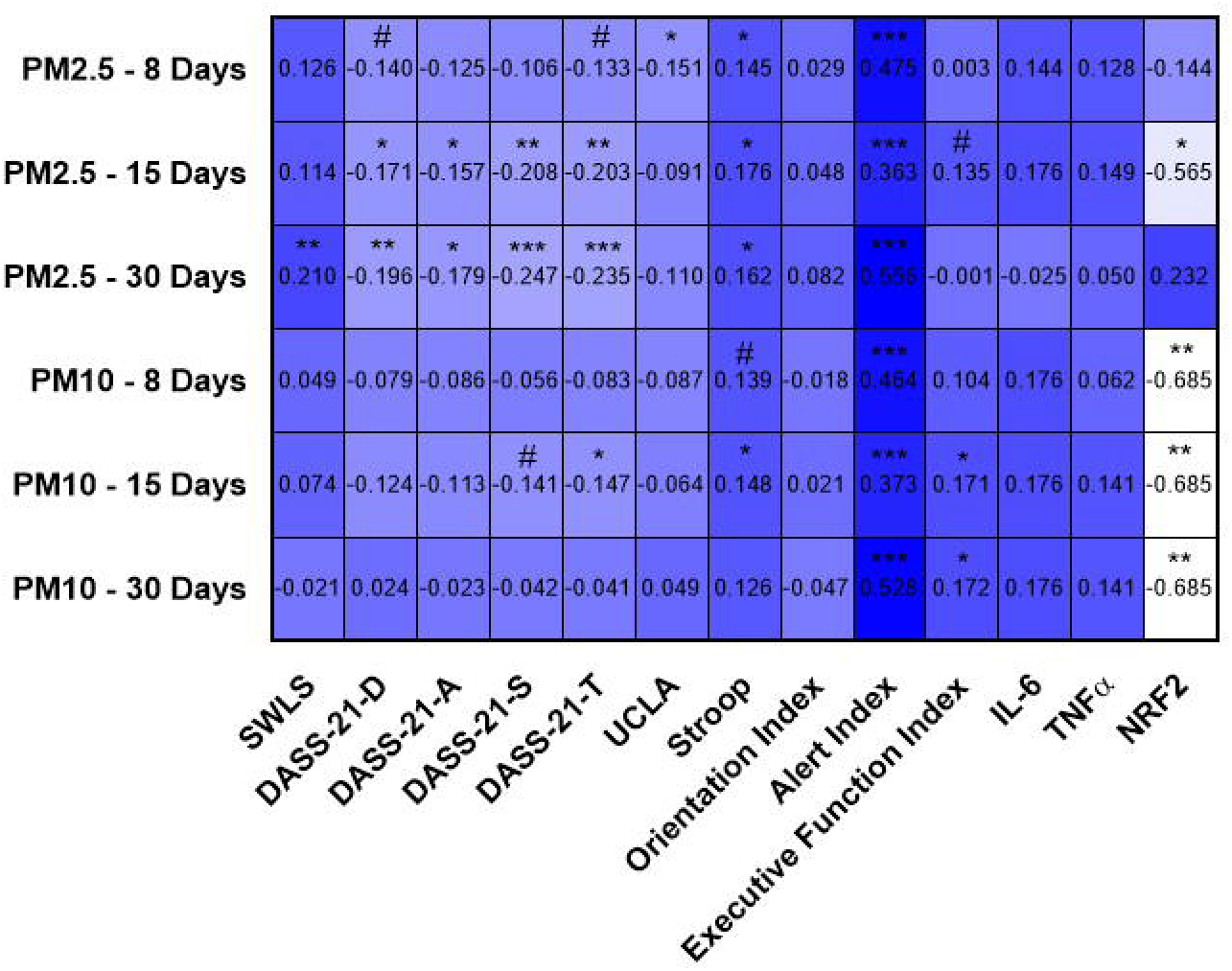
Heatmap representation of Spearman correlations between molecules and exposure times with some variables. Only significant comparisons and tendency correlations are depicted. Only significant correlations are depicted. The number in each quadrant represents the calculated Rho number. Significance levels are depicted with the following symbols: ***p <.001, **p <.01, *p <.05, # p <.07.

Second, positive correlations were detected between PM2.5 exposure for all time periods analyzed with behavioral outcomes [8-days, Stroop (p =.05, ρ =.145); ANT, Alert Index (p <.001, ρ =.475); 15-days, Stroop (p =.017, ρ =.176); ANT, Alert Index (p <.001, ρ =.363), Executive Functioning (p =.069, ρ =.135); 30-days, Stroop (p =.028, ρ =.162); ANT, Alert Index (p <.001, ρ =.556)]. Furthermore, positive correlations were present in PM2.5 exposure for 30 days with life satisfaction (p =.004; ρ =.210).

Third, PM10 exposure windows did not achieve the same correlations as PM2.5. Only PM10 exposure time for 15-days achieve anecdotal correlations with stress (p =.055, ρ = -.141) and significant correlation with total punctuation (p =.046, ρ = -.147). On the contrary, and same as with PM2.5, positive correlations were detected with behavioral outcomes for 15-days [Stroop (p =.045, ρ =.148); ANT, Alert Index (p <.001, ρ =.373), Executive Function Index (p =.021, ρ =.171)]. Higher PM concentrations are related with slower reaction time (ANT) or higher interference (Stroop)

Lastly, regarding biochemical outcomes, we found several negative correlations between PM10 exposure windows and NRF2 concentrations (all p’s =.004, ρ = -.685), while 15-days exposure of PM2.5 is also negatively correlated (p =.025, ρ = -.565). Higher PM exposure is related to lower NRF2 concentrations.

Therefore, it is noteworthy the difference in the number of correlations between PM2.5 and PM10 with our outcomes. PM2.5 is more related with our variables, achieving more correlations with higher exposure time. Also, seems important the relationship between all exposure windows and all molecules with the Alert Index in the ANT task.

### 3.4. Relationship between in silico MPPD simulated absorption PM10 and PM2.5 concentration and emotional variables

All the linear regression analyses were performed with MPPD-simulated absorbed data from three time-period windows exposure: 8-, 15-, 30-days prior data collection and all the cognitive/behavioral data.

Starting with cognitive/emotional variables, PM2.5 did not predict life satisfaction outcomes [8-days, p =.149; 15-days (F_(2, 185)_ = 2.444; p =.090; R^2^ =.026); 30-days (F_(2, 185)_ = 2.736, p =.068; R^2^ =.029)]. However, in both time exposure, PM coefficient achieved significant levels [15-days, t = 2.200, p =.029, R^2^ =.020; 30-days, t = 2.329, p =.021, R^2^ =.024; Figure 5d-5f] predicting life satisfaction. Higher PM2.5 exposure predicts better life quality punctuations. Regarding PM10, none of the analyzed windows achieved neither significant levels nor anecdotal.

**Figure 5.**
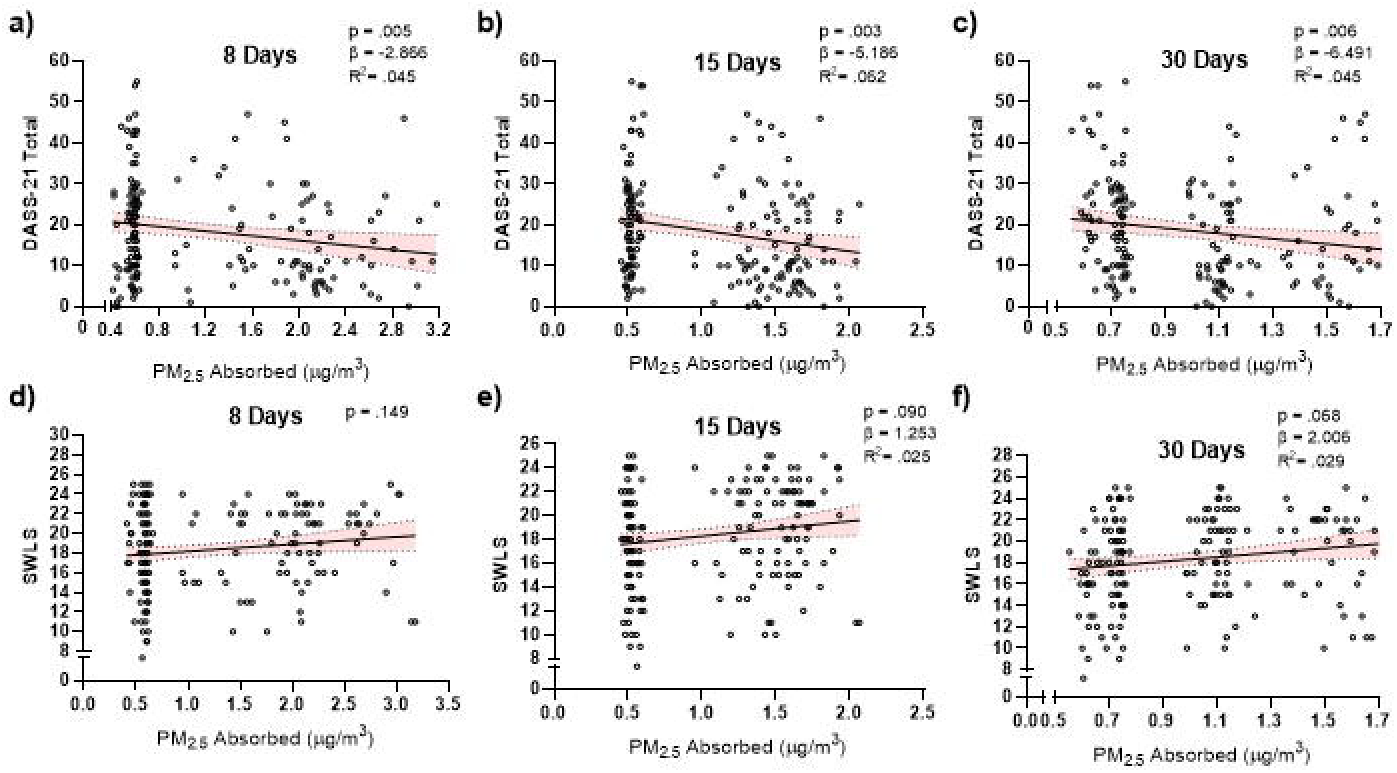
Linear relationships between PM2.5 Absorbed derived from the MPPD model and different cognitive outcomes. lmages a-c depicts its relationship with DASS-21 total score. lmages d-f shows its relationship with SWLS (life satisfaction).

The same models were calculated for DASS-21 total punctuation and its subsections. All PM2.5 exposure windows achieved significant levels for total punctuation [8-days (F_(2, 184)_ = 5.361, p =.005, R^2^ =.045; 15-days (F_(2, 184)_ = 6.049, p =.003, R^2^ =.062); 30-days (F_(2, 184)_ = 5.315, p =.006, R^2^ =.045); Figure 5a-5c]. Sex reached significant levels at 8-days (t = 2.014, p =.045) and at 30-days (t = 2.126, p =.035). PM2.5 concentrations were significant in all exposure windows (p <.05). All PM10 models analyzed were able to predict total DASS-21 punctuations, but in all models, only sex achieved significant levels while PM10 concentrations remained not-significant [8-days (F_(2, 184)_ = 3.289, p =.040, R^2^ =.024); 15-days (F_(2, 184)_ = 2.991, p =.053, R^2^ =.021); 30-days (F_(2, 184)_ =.025, p =.037, R^2^ =.036); Data not shown]. Higher exposure to PM is related with lower punctuations in total DASS-21 punctuation. The depression subscale follows the analysis. First, regarding PM2.5 exposure, only 15-day exposure reached significance (F_(2, 184)_ = 3.111, p =.047, R^2^ =.033), while the 8-, and 30-days did not predict the subscale punctuations (Figure 6a-6c). Higher PM2.5 exposure is linked to lower depression scores in the DASS-21 subscale. Neither of the PM10 exposure windows predicted the DASS-21 depression subscale outcomes. In terms of anxiety subscale, 8-, and 30-days predict anxiety outcomes, while the 15-days remain at a trend level. Only sex reached significance in those models [8-day (F_(2, 184)_ = 4.740, p =.010, R^2^ =.050); 15-days (F_(2, 184)_ = 4.194, p =.017, R^2^ =.044); 30-days (F_(2, 184)_ = 4.216, p =.016, R^2^ =.044); Figure 6d-6f]]. Same pattern was found for PM10 [8-days (F_(2, 184)_ = 3.277, p =.040, R^2^ =.035); 15-days (F_(2, 184)_ = 2.794, p =.064, R^2^ =.030); 30-days (F_(2, 184)_ = 3.253, p =.041, R^2^ =.035); data not shown]. Lastly, in terms of the stress subscale, all exposure windows achieved significant levels for PM2.5 [8-days (F_(2, 184)_ = 6.802, p =.001, R^2^ =.070); 15-days (F _(2, 184)_= 8.609 p <.001, R^2^ =.076); 30-days (F_(2, 184)_ = 8.039, p <.001, R^2^ =.071). In all models sex and PM exposure time reached significant levels [8-days (PM2.5; t = -1.982, p =.049; sex; t = 2.749, p =.007); 15-days (PM2.5; t = -2.714, p =.007; sex; t = 2.313, p =.022); 30-days (PM2.5; t = -2.506, p =.013; sex; t = 2.811, p =.005); Figure 6g-i]. Same pattern as before was detected for PM10 [8-day (F_(2, 184)_ = 5.067, p =.007, R^2^ =.053); 15-days (F_(2, 184)_ = 5.184, p =.006, R^2^ =.054); 30-days (F_(2, 184)_ = 4.853, p =.009, R^2^ =.051); data not shown]. Sex was the only significant variable in all exposure windows (p <.05). Linear regression showed that higher PM exposure leads to lower stress punctuation in the DASS-21 subscale; furthermore, women had higher punctuations in this subscale.

**Figure 6.**
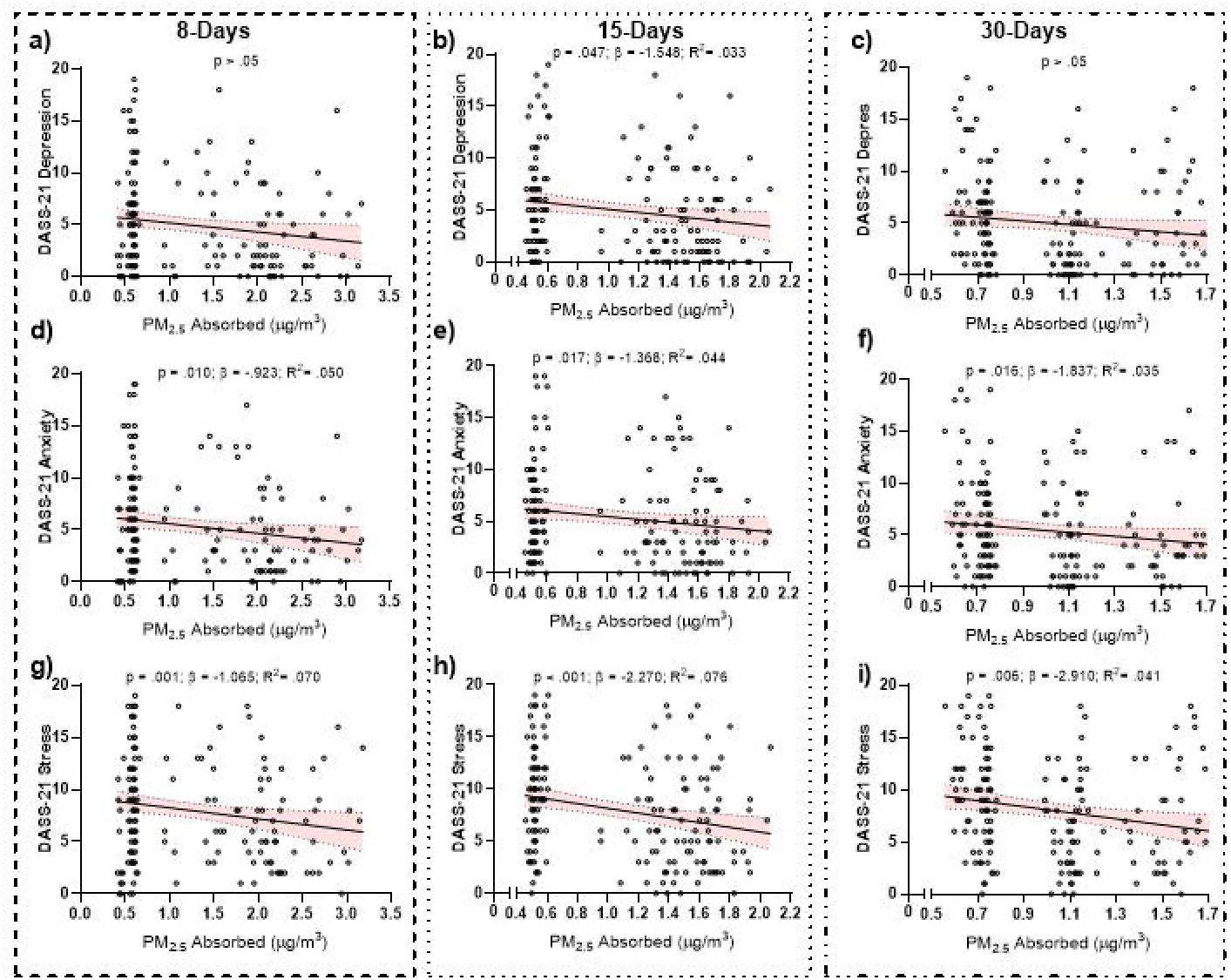
Linear relationships between PM2.5 Absorbed derived from the MPPO model and DASS-21 subscales. Figure a-c represent depression, figures d-f correspond to anxiety and figures g-l correspond to stress subscales. AII graphs depict individual data.

The same analyses were performed with loneliness. No model achieved significant levels, neither those with PM2.5 nor with PM10 and loneliness (all p’s >.05) The same result was found for PM10. Lastly, in terms of impulsivity, no time exposure neither molecule was able to predict either of the BIS-11 outcomes (total, non-planned, motor and cognitive impulsivity).

### 3.5. Relationship between in silico MPPD simulated absorption PM10 and PM2.5 and cognitive variables

Regarding the Orientation Index (time spent shifting spatial attention) calculated from valid and invalid cues in the ANT, neither PM molecules neither exposure time windows achieved significance as predictors of orientation index (p >.05). Same results were detected from the Alert Index (time spent increasing alert readiness) calculated with the alerted cue and non-cue. None of the models tested achieved significant levels (p >.05). However, the results found for Executive Index (time spent resolving stimulus conflict) are different from the previous indexes. Neither of the exposure windows for PM_2.5_ achieved significance (p >.05). PM_10_ exposure window for 8-days did not achieve significance (p =.192); nevertheless, PM_10_ exposure for 15-days achieved moderate fit (F_(1, 181)_ = 4.573; p =.034; R^2^ =.025). Also, PM_10_ exposure window for 30-days achieved significant levels (F_(1, 181)_ = 4.153; p =.043; R^2^ =.023; Figure 7a-c). The best predictor is the 15-day period, which explained 1.9% of the variance. In terms of 15-day exposure, no significance was detected for Congruent-(p =.360), Incongruent-(p =.572) nor Neutral-trials (p =.326). For the 30-day period, no significance was detected for Congruent-(p =.764), Incongruent-(p =.149), nor Neutral-trials (p =.527).

**Figure 7.**
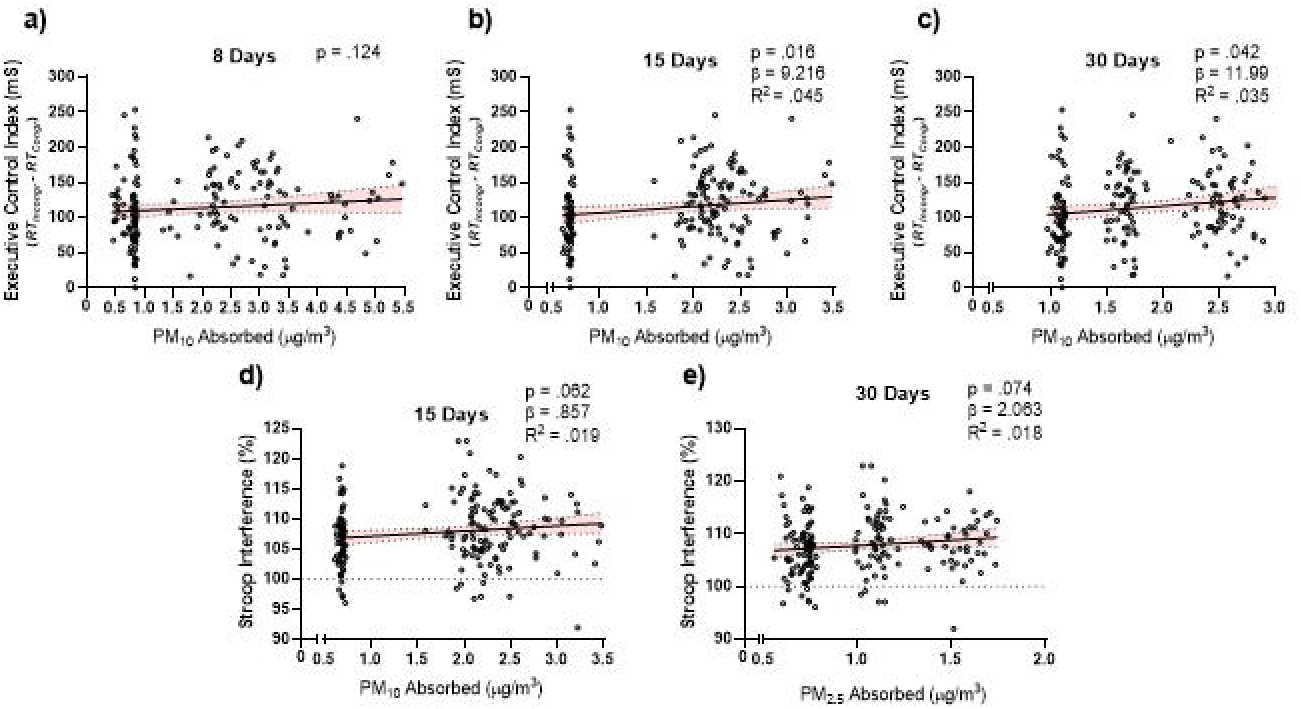
Graphical representation of linear regression between time exposure periods and cognitive variables. Upper row (images a to c) represents ANT Executive Control Index, while the lower row images represent Stroop outcomes.

Regarding Stroop-outcomes, no significant outcomes were present for PM10, while 15-days PM10 exposure remained anecdotal (F_(1, 183)_ = 3.533; p =.062; R^2^ =.019). In terms of PM2.5, the 15-day exposure window achieved significant levels (F_(1, 183)_ = 4.430; p =.037; R^2^ =.024), while the other exposure windows did not achieve significant levels (Figure 7d-e).

### 3.6. Relationship between in silico MPPD simulated absorption PM10 and PM2.5 concentration and ELISA outcomes

Due to the presence of heteroscedasticity and deviations from normality in the residuals, a logarithmic transformation was applied to the ELISAs variables prior to regression modeling. Regarding IL-6 data, no model reached evidence (p >.05). Same results were found for Klotho protein concentration (p >.05), and for TNF-α concentration (p >.05).

On the contrary, several models reached significant levels regarding NRF2a protein concentration. In terms of PM2.5 exposure periods, only the 15-day exposure period reached significant levels [8-days (p >.05); 15-days (F_(2, 15)_ = 7.527; p =.007; R^2^ =.465); 30-days (F_(2, 15)_ = 1.017; p =.389; R^2^ =.002); Figure 8a-c]. On the contrary, all the exposure windows of PM10 exposure reached significant levels [8-days (F_(2, 15)_ = 5.832; p =.016; R^2^ =.392); 15-days (F_(2, 15)_ = 5.709; p =.017; R^2^ =.468); 30-days (F_(2, 15)_ = 6.034; p =.014; R^2^ =.481); Figure 8d-f]. PM and NRF2a dynamics are complex, showing that being exposed to PM for higher concentrations is link with lower NRF2a, but this dynamic is not present for PM2.5.

**Figure 8.**
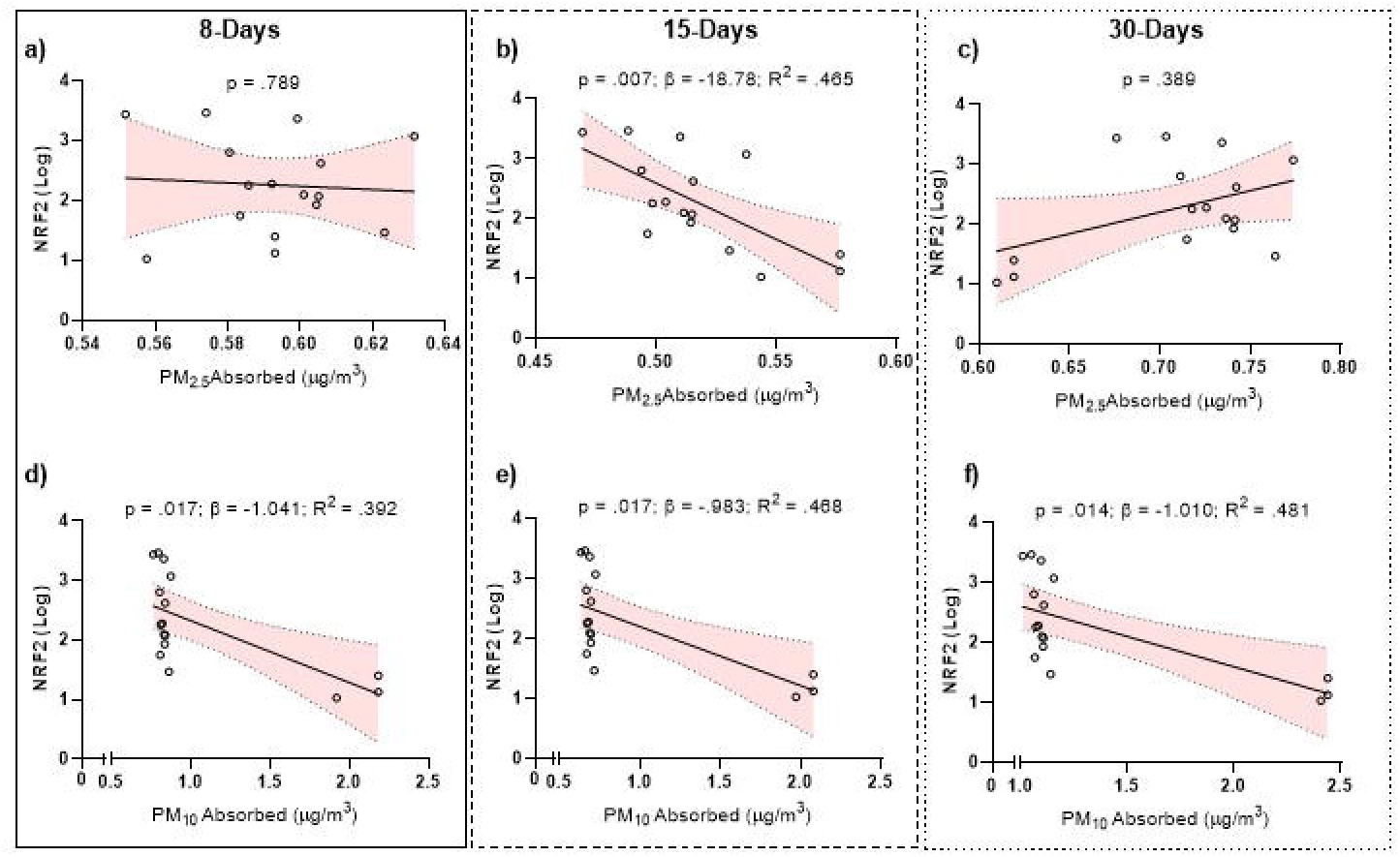
Graphical representation of the Linear Regression analysis between PM2.5 and PM10 exposure with NRF2a protein concentration analyzed with ELISA. Upper graphs (a-c) correspond to in silico PM2.5 absorption calculation; lower graphs (d-f) correspond to in silico PM10 absorption calculation. Each graph represents individual data.

## 4. Discussion

The objective of the present study was to examine the effects of short-term (8 days), mid-term (15 days), and long-term (30 days) exposure to in silico PM10 and PM2.5 exposure on cognitive functioning, with a particular focus on attention and cognitive interference control. Additionally, systemic biological markers were incorporated to explore potential indirect inflammatory and oxidative stress pathways associated with these behavioral outcomes. Individual PM exposure concentrations were estimated using the MPPD model, allowing for a personalized assessment of inhaled particle deposition. To our knowledge, this is the first study to use the MPPD model to simulate PM exposure in human subjects and to apply these data to predict cognitive and emotional outcomes. Furthermore, no previous studies have examined these effects in healthy young individuals in three different cities, which represent markedly distinct environments and therefore capture the heterogeneity of contemporary Western contexts.

We observed that PM10 and PM2.5 exposure across 8-, 15-, and 30-day periods was associated with lower scores in total psychological symptoms and higher reported quality of life. In addition, exposure to PM10 over 15 and 30 days was related to poorer executive control and greater interference effects in behavioral performance. Furthermore, PM10 exposure was significantly associated with serum NRF2 levels, supporting a potential relationship between particulate matter exposure and oxidative stress regulation.

A key methodological strength of this study lies in the use of the MPPD model to estimate individualized inhaled particle deposition (Junaidi et al., 2025). Unlike previous epidemiological studies that rely primarily on ambient concentration measures, the MPPD approach provides a biologically grounded estimation of deposited dose within the respiratory tract (Junaidi et al., 2025; Choi et al., 2025). Ambient concentrations do not necessarily reflect internal exposure, as actual deposited dose depends on multiple physiological and particle-specific factors. By modeling particle deposition, this approach approximates the biologically effective dose reaching the lower respiratory tract, which may be more directly associated with systemic responses (Alexis et al., 2006). Moreover, traditional exposure metrics based on area-level pollution may introduce non-differential exposure misclassification, potentially attenuating observed associations (Yu et al., 2024). Individualized deposition modeling may partially mitigate this limitation and enhance exposure precision. This improved exposure characterization may help explain the detection of subtle associations between particulate matter exposure and attentional functioning observed in the present study.

Most evidence in environmental toxicology remains grounded in preclinical models (rodents, fish, or nematodes). In human populations, the majority of studies have relied on cohort designs to examine the relationship between particulate matter exposure and cognitive functioning. For example, Qi et al. (2024) analyzed three follow-up waves within the Chinese Health and Retirement Longitudinal Study (CHARLS; 2000–2018) and reported that long-term exposure to higher concentrations of PM2.5 was associated with an increased risk of cognitive impairment (Hazard Ratio [HR] = 1.19). However, that study focused on individuals older than 65 years, a population already within the age range at risk for age-related cognitive decline. Our findings extend this literature by suggesting that pollution-related cognitive alterations may also be detectable outside advanced aging populations.

Additional analyses using the CHARLS database (Yao, Wang, & Xiang, 2022) reported that PM1, PM2.5, and PM10 were associated with cognitive decline in middle-aged adults, with effects moderated by sex, region, and lifestyle factors. These findings highlight the potential role of sociodemographic and behavioral variables in shaping vulnerability to air pollution exposure, an issue that warrants further exploration.

A growing body of research has also examined cognitive outcomes in children. Jalaludin et al. (2022) reported a small positive association between PM2.5 exposure and cognitive scores in Indonesian regions affected by forest fires, whereas Sukumaran et al. (2024) found negative associations between PM2.5 components and multiple neurocognitive domains (general ability, memory, executive function) in U.S. children aged 9–10 years. These heterogeneous findings suggest that the impact of particulate matter may vary according to exposure context, developmental stage, and pollutant composition. A systematic review by Donzelli et al. (2019) reported inconsistent evidence linking particulate matter to ADHD but noted a clearer association with attentional and behavioral problems. Similarly, Saenen et al. (2016) observed impaired selective attention in Belgian children, and Siddique et al. (2010) reported a positive association between PM10 exposure and ADHD symptoms in Indian youth. Same as with epidemiological data, Faherty et al. (2025) performed an experiment where the PM concentrations were modified using candles; they found an increased reaction time in the executive control network after being exposed to PM for 1 hour.

In Spain, Sunyer et al. (2015) reported that children attending highly polluted schools in Barcelona exhibited reduced cognitive development growth compared to peers in less polluted schools. In contrast, Gignac et al. (2021) found no significant improvements in attentional outcomes following air filtration interventions in adolescents. To our knowledge, however, limited evidence has examined these associations in youth populations using individualized exposure estimation approaches.

Collectively, these findings indicate that PM exposure is associated with measurable behavioral and cognitive alterations. Our results are broadly consistent with previous literature linking particulate matter exposure to impaired cognitive performance. To our knowledge, this is the first study to detect altered cognitive performance in youth using individualized exposure estimation without experimentally manipulating ambient PM concentrations. Faherty et al. (2025) were the first to experimentally modify PM concentrations and demonstrate attentional alterations following short-term exposure. Our findings extend their results by suggesting that similar cognitive effects may be detectable under real-world exposure conditions. However, several methodological considerations warrant further discussion. First, neither study comprehensively adjusted for potential confounding variables, whereas large-scale epidemiological investigations typically incorporate extensive covariate control. Second, Faherty et al. (2025) employed controlled exposure manipulation, while our design relied on naturally occurring exposure variability, which limits causal inference. Third, both approaches are unable to characterize the specific chemical composition of particulate matter, which may critically influence biological and cognitive outcomes. Neuroimaging research provides additional context for these findings. Previous studies have reported alterations in default mode network functioning in children exposed to higher pollution levels (Pujol et al., 2016), with more recent evidence indicating disrupted network dynamics associated with PM2.5 exposure (Zundel et al., 2024). Taken together, these data suggest that the cognitive effects observed in the present study may reflect subtle pollution-related alterations in large-scale brain network organization.

The influence of sex as a covariate appears only in the DASS-21 anxiety and stress subscales, while the remaining variables are minimally or not affected at all. This pattern is consistent with the well-established literature showing multiple sex-related differences in stress responses. Stress reactivity is one of the domains with the clearest sex differences (Verma et al., 2011). The DASS-21 is a self-report instrument, and women typically report higher levels of subjective distress in response to stressors (Childs et al., 2010; Kelly et al., 2008), whereas men tend to show stronger physiological stress reactivity. Moreover, men and women often engage in qualitatively different coping strategies when facing stress (Nolen-Hoeksma, 2012).

Regarding biochemical pathways, evidence linking air pollution, systemic inflammation, and cognitive functioning in humans remains limited. Most mechanistic insights derive from preclinical models (Ehsanifar et al., 2022; Ruiz-Sobremazas et al., 2023). Some human studies suggest associations between pollutant exposure and inflammatory markers. For instance, Du et al. (2022), using CHARLS data, reported that solid fuel use was associated with higher depression risk, poorer cognitive functioning, and elevated white blood cell counts. However, controlled exposure studies in healthy adults have not consistently demonstrated acute changes in classical pro-inflammatory markers following diesel exhaust exposure (Cliff et al., 2016). In line with these findings, our results did not show significant associations between particulate matter exposure and traditional inflammatory markers.

Notably, however, we observed a significant association between PM10 exposure and NRF2 levels. NRF2 is primarily involved in oxidative stress regulation rather than classical inflammatory signaling. Preclinical evidence indicates that PM2.5 exposure can alter NRF2 activity (Mei et al., 2024), and interactions between NRF2 and NF-κB pathways are known to modulate oxidative stress and inflammatory responses (Zhang et al., 2021; Gao et al., 2022). The interplay between these pathways is complex, as NF-κB activity may suppress NRF2 signaling, while NRF2 deficiency can enhance NF-κB–mediated inflammatory responses. Future studies should simultaneously assess NRF2 and NF-κB activity to better characterize the oxidative–inflammatory balance associated with particulate matter exposure. There is not enough information about the impact of PM on inflammatory or oxidative stress markers in humans.

The present study is not without limitations. First, the sample size was relatively modest, which may have limited statistical power to detect small associations and to explore these relationships in greater detail. Although many epidemiological studies rely on substantially larger samples, the present work focused on environmental toxicological mechanisms, which involve more intensive individual-level exposure and biomarker assessments. Second, the range of particulate matter concentrations was geographically restricted. Including participants from a broader range of Spanish cities, as well as rural areas, would allow greater variability in exposure levels and enhance the generalizability of the findings. Third, future studies should incorporate additional covariates to better account for potential confounding factors, including lifestyle, socioeconomic status, and pre-existing health conditions. Fourth, serum clotting time was not standardized, which may have introduced additional variability in some biochemical measures. Finally, experimental or quasi-experimental designs that manipulate or more precisely control exposure periods, such as approaches integrating prior exposure modeling using MPPD, may further clarify temporal dynamics and dose-response relationships.

Further research is needed to continue exploring the long-term consequences of PM exposure during early life, particularly to determine how such exposure may influence health outcomes later in adulthood and aging. Understanding whether early exposure predisposes individuals to future cognitive, emotional, or oxidative stress–related alterations remain a key research priority. Longitudinal studies will be essential to clarify potential delayed effects and to establish causal pathways across the lifespan.

Nevertheless, the present study provides novel evidence in this field, as it uniquely applies the MPPD model to simulate PM exposure in young healthy individuals and examines its association with cognitive, emotional, and biological markers, revealing meaningful relationships between environmental pollution variables and executive functioning, as well as an interesting pattern involving NRF2. Importantly, these findings emerge from three cities with completely distinct geographical characteristics, offering a set of contexts that are highly representative of Western heterogeneity.

## Acknowledgements

The present work is part of the grant reference: PID2020-113812RA-C33 funded by the Spanish Government (Ministry of Science and Innovation; MCIN/AEI/10.13039/501100011033) (C.L-G), PROY_S12_24 by Government of Aragon, R&D&I in priority and multidisciplinary research lines (C.L-G) and by European Union, Next Generation EU (Investigo) (D. R-S; C.L-G).

Diego Ruiz-Sobremazas (Methodology, Software, Validation, Formal analysis, Investigation, Data Curation, Writing, Visualization), Blanca Cativiela-Campos (Investigation, Data Curation, Writing); María Cadalso (Investigation, Data Curation, Writing); Angel Barrasa (Methodology, Investigation, Data Curation, Writing), Pilar Catalán-Edo (Methodology, Investigation, Data Curation, Resources, Writing, Visualization); Cristian Perez-Fernandez (Investigation, Data Curation, Writing); Beatriz Ferrer (Investigation, Formal Analysis, Supervision); Fernando Sánchez-Santed (Investigation, Data Curation, Writing); Teresa Colomina (Investigation, Data Curation, Writing); Caridad López-Granero (Methodology, Software, Validation, Formal analysis, Investigation, Data Curation, Writing, Visualization, Funding Acquisition).

